# Whole genome assembly of *Culex tarsalis*

**DOI:** 10.1101/2020.02.17.951855

**Authors:** Bradley J. Main, Matteo Marcantonio, J. Spencer Johnston, Jason L. Rasgon, C. Titus Brown, Christopher M. Barker

## Abstract

The mosquito, *Culex tarsalis*, is a key vector in the western United States due to its role in transmission of zoonotic arboviruses that affect human health. Extensive research has been conducted on *Cx. tarsalis* ecology, feeding behavior, vector competence, autogeny, diapause, genetics, and insecticide resistance. Population genetic analyses in the western U.S. have identified at least three genetic clusters that are geographically distinct. Salivary gland-specific gene expression has also revealed genes involved in blood feeding. However, genetic studies of this mosquito have been hindered by the lack of a reference genome. To facilitate genomic studies in *Cx. tarsalis*, we have assembled and annotated a reference genome (CtarK1) based on PacBio HiFi reads from a single male. Using the *Cx. tarsalis* transcriptome and protein sequences from *Culex quinquefasciatus*, approximately 17,456 protein-coding genes, including the *para* insecticide resistance gene, were annotated in the CtarK1 genome. Genome completeness was assessed using the Benchmarking Universal Single-Copy Orthologs (BUSCO) tool, which identified 84.8% of the 2799 Dipteran BUSCO genes. The CtarK1 assembly is 790Mb with an N50 of 58kb. Using full mitochondrial genome alignments with other sequenced mosquito genomes we present a Bayesian phylogeny, which estimates that the divergence of *Cx. tarsalis* from *Culex quinquefasciatus*, the most closely related mosquito species with a genome, occurred 15.8-22.2 million years ago.

## Introduction

The mosquito, *Culex tarsalis*, is one of the most important vectors in the western United States due to its capacity to transmit several arboviruses that cause disease in humans and horses. In agricultural areas, it is the principal vector for West Nile virus (Goddard *et al*. 2002; Reisen 2013), and such areas have the highest incidence of West Nile virus disease. Extensive research has been done on *Cx. tarsalis* ecology (Reisen 2012), feeding behavior (Thiemann *et al*. 2012; Reisen *et al*. 2013), vector competence (Kramer *et al*. 1981; Reisen *et al*. 2014), autogeny (Spadoni *et al*. 1974; Reisen 1995), diapause (Reisen 1986; Buth *et al*. 1990; Reisen *et al*. 1995), and insecticide resistance (Ziegler *et al*. 1987). A genetic linkage map has also been developed (Venkatesan *et al*. 2009) and three genetically distinct populations have been described in the western US: the Pacific, Sonoran, and Midwest genetic clusters (Venkatesan and Rasgon 2010). However, the lack of a published reference genome has likely limited in-depth genomic analyses in *Cx. tarsalis*.

Here we describe the first genome assembly for *Cx. tarsalis*. The assembly is based on PacBio HiFi reads from a single adult male and 10X genomics data was used for assembly of the mitochondrial genome. We used DNA from a male as input so that the dominant sex locus could be identified as well as the female sequence at that region. This resource will facilitate the characterization of sex determination in the system, future landscape genetics/genomics, and additional phylogenetic studies among mosquito species.

## Methods

### Mosquitoes

All *Cx. tarsalis* used for the genome assembly were sampled from the Kern National Wildlife Refuge (KNWR) colony, which was established in 2002 from mosquitoes collected at the Kern National Wildlife Refuge (35.7458°N, 118.6179°W), in Kern County, CA, USA. For PacBio sequencing, high molecular weight (HMW) DNA was extracted at the UC Berkeley DNA Sequencing Facility from a single male adult. For 10X genomics library preparation, HMW DNA was extracted by the UC Davis DNA Technologies Core from two late-eclosing, relatively large pupae (likely female). Two pupae were used for 10X because DNA yields were too low from a single individual. In addition, DNA from a single adult male was used to make a Nextera library (Illumina) for standard genome sequencing with paired-end 75bp reads.

### PacBio genome assembly

PacBio sequencing was performed on a Sequel II SMRT cell at the UC Berkeley sequencing core. Circular consensus sequences (CCS) were then generated and filtered with high stringency to get HiFi CCS reads. We generated an initial genome assembly using Canu v1.9 with the following settings: genomeSize=0.875g, useGrid=false. The genome coverage was low, but because these were HiFi reads, we reduced the standard minimum coverage thresholds with the following options: stopOnLowCoverage=1, contigFilter=“2 0 1.0 0.5 0”. The PacBio mitochondrial contig appeared to be two full mitochondrial genomes stuck end to end. As a result, we removed this contig and replaced it with the full mitochondrial genome generated from the 10X assembly. Then we performed genome polishing using racon (v1.4.3) and 75bp Nextera reads (Illumina) from a single adult male from the KNWR colony. Additional SNPs in the reference sequence compared to multiple sequenced KNWR individuals were removed by extracting the consensus sequence from a single male KNWR library mapped to the reference using bcftools. Annotations were performed using Maker (v.2.31.10) with the *Culex tarsalis* transcriptome (Ribeiro *et al*. 2018) and *Cx. quinquefasciatus* protein sequences (CpipJ2.4). Repeat masking was performed in parallel using the maker annotation pipeline. As input, we used the standard RepBase database (RepBaseRepeatMaskerEdition-20170127) and a *Cx. tarsalis*-specific repeat library that was generated using RepeatModeler (v1.0.11). The mitochondrial genome and the para gene were re-annotated using Geneious software and the *Cx. quinquefasciatus* mitochondrial genome (NC_014574) as a reference. To calculate genome statistics, including the N50, we used Quast (v5.0.2).

### 10X linked-read genome assembly

The HMW DNA extraction, 10X Genomics Chromium sequencing library, and Illumina sequencing was performed at the UC Davis DNA Technologies Core. Raw linked reads were assembled using the Supernova 2.0.1 software (10X Genomics) on an AWS instance with 480Gb on RAM and 64 logical cores. To generate a fasta formatted reference sequence, we used supernova mkoutput with --style=pseudohap. This option arbitrarily selects haplotypes across the genome resulting in one pseudo-haploid assembly composed of a mosaic of paternal and maternal haplotype stretches.

### Phylogenetic analyses

The mitochondrial analysis included 15,203bp of mitochondrial sequence from each species: *Cx. tarsalis* (CtarK1, this study), *Cx. quinquefasciatus* (NC_014574), *Anopheles gambiae* (NC_002084), *Anopheles coluzzii* (NC_028215), *Aedes aegypti* (NC_035159.1), *Aedes albopictus* (NC_006817), and *Drosophila melanogaster* (NC_024511). A multiple sequence alignment was performed using MUSCLE (v3.8.425; (Edgar 2004)) and then the phylogenetic analysis was performed using the Bayesian Evolutionary Analysis by Sampling Trees software (BEAST) v 2.6.2 (Drummond and Rambaut 2007). In order to determine the best combination of substitution and clock models, model selection was performed through Generalised stepping-stone sampling (GSS) using BEAST Path Finder tool (1 M iterations and 100,000 pre-burnin). The model with the lowest marginal log-likelihood had Generalised time-reversible substitutions (GTR, with 4 category count for the Gamma site model), relaxed log-normal clock, and Yule tree model. In addition, to improve the estimated divergence time for the *Culex* clade, we used a Yule calibrated tree model using the following previously published tree calibration constraints: 71 (44.3, 107.5) million years ago (MYA) for *Aedes aegypti* and *Aedes albopictus*, 179 (148.0, 216.7) MYA between *Culex* and *Aedes*, 217 (180.8, 256.9) MYA between *Culicinae* and *Anopheles*, and 260 (238.5, 295.5) MYA between *Drosophila* and *Culicidae* (Chen *et al*. 2015). All clades with priors were considered monophyletic. The final phylogenetic tree was generated running 100 M iterations with 10 M burn-in saving samples every 1000 steps in order to obtain a representative sample (i.e. estimated sample size [ESS] higher than 200 for all important model parameters) for the posterior distribution of model parameters. The maximum clade credibility tree with average node heights was generated in TreeAnnotator, by removing the initial 10% fraction of the chain.

### Genome size estimate

Genome size was estimated as described in Johnston, Bernardini & Hjelmen 2019 (Johnston *et al*. 2019), and is based on the fluorescence scored using a Partex CX flow cytometer that was equipped with green laser excitation. Briefly, the head of a single *Cx. tarsalis* mosquito was combined with the head of a *Drosophila virilis* standard (1C = 328 Mbp) in 1 ml of cold Galbraith buffer and ground with 15 strokes of the “A” pestle in a 2ml Kontes Dounce. The nuclei released were filtered through a 45 U nylon mesh, stained for 1 hour in the cold and dark using 25 ug/ml propidium iodide. The total mean fluorescence of the 2C (diploid) peaks from the sample and standard was measured as a mean channel number using the software supplied with the Partec Cx. The 1C amount of DNA in the mosquito was estimated as (mean channel number of the 2C peak of *Cx. tarsalis* / mean channel number of the 2C *D. virilis* peak) X 328 Mbp. More than 2,000 nuclei were scored under each 2C peak and the CV of the peaks were all < 2.5).

## Results and Discussion

### Genome assembly

The PacBio long-read library generated from a single adult male *Cx. tarsalis* was sequenced on the Sequel II platform and yielded 5.7M raw reads and 988,512 ccs HiFi reads. The HiFi reads were used to prepare a draft assembly using Canu with reduced filtering thresholds due to the lower error rate in these reads (see methods). This resulted in 19,994 contigs and a total genome size of 789,669,425bp (Quast v5.0.2). The N50 was 57,901bp, the GC content was 36%, and the largest contig was 753,184bp (Table 1). To assess the genome completeness of the CtarK1 PacBio assembly, we searched for the presence of a set of 2799 Dipteran Benchmarking Universal Single-Copy Orthologs (BUSCO; (Zdobnov *et al*. 2017; Waterhouse *et al*. 2018). We detected 79% (2219/2799) as complete single copy genes, 8% (227/2799) as complete and duplicated, 5% (153/2799) were fragmented, and 15.2% (427/2799) were missing.

**Table 1.** Assembly statistics.

| Assembly | Genes | Contig # | Median contig (N50) | Total length |
| --- | --- | --- | --- | --- |
| *Cx. quinquefasciatus* (CpipJ2.4) | 19,793 | 3,172 | 486,756 bp | ~579Mb |
| *Cx. tarsalis* (CtarK1) | 17,456 | 19,994 | 57,901 bp | ~790Mb |

Using flow cytometry, we estimated the complete haploid genome size of *Cx. tarsalis* to be 890Mb. This genome size estimate and the BUSCO scores from our 790Mb PacBio assembly indicate that this initial assembly is approximately 85-90% complete. Additional long-read sequencing from single individual mosquito input and genetic scaffolding (e.g. with Hi-C) is needed to make CtarK1 comparable in quality to other model species.

For the mitochondrial genome, we used a 10X chromium library that was sequenced on Illumina’s Novaseq platform, yielding approximately 507 million clusters (paired-end reads) passing filter. Based on several trial assemblies, we downsampled the total read input to 350 million paired-end reads to yield approximately 56x coverage. A contig containing the complete mitochondrial genome was then identified, and redundant sequence (due to circular genome) was trimmed at each end of the contig. While we included the 10X mitochondrial genome in our CtarK1 assembly and we used 10X contigs to make scaffolds at the sex locus (described later), we avoided making a consensus genome between PacBio and 10x because of concern over high levels of redundancy in 10X assembly. For example, the final 10x assembly size (1.5Gbp) was nearly twice the expected size (890Mb). This was likely a result of inputting multiple pupae, which resulted in multiple haplotypes in the assembly.

### Genome annotation

A total of 14,726 complete genes were identified by the maker annotation pipeline (see methods). An additional 2,730 gene orthologs have support from protein2genome annotations with an arbitrary overlap threshold of 80% or less. Thus, we identified approximately 17,456 genes in CtarK1. While this is substantially fewer than the number for *Cx. quinquefasciatus*, direct comparisons should not be made until BUSCO scores are similar between the assemblies.

### Pyrethroid resistance locus

The emergence of resistance to pyrethroid insecticides in *Cx. tarsalis* is an important issue that needs to be considered and managed for vector control efforts. The best-studied genetic mutation underlying pyrethroid resistance is a L1014S or L1014F mutation that occurs on exon 6 (see asterisk on Figure 1) of the voltage-gated sodium channel gene (*para*). This mutation is referred to as knockdown resistance (*kdr*).

**Figure 1.**
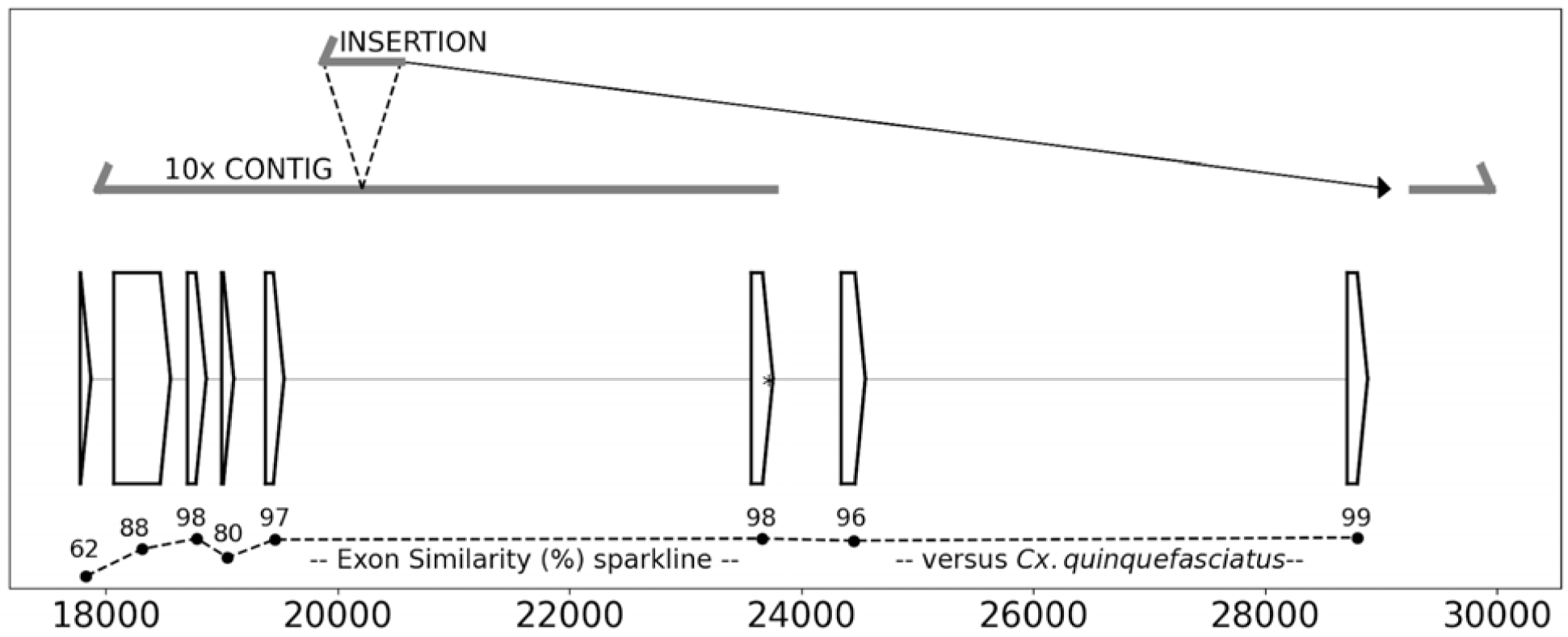
PacBio contig spans the entire *para* gene. Annotations of the insecticide resistance gene *para* on tig00032677 of the CtarK1, including coding sequences (open boxes), the location of *kdr* in exon 6 (*), and the percent similarity of each coding region compared to *Cx. quinquefasciatus* (--). In contrast to PacBio, the corresponding 10X Genomics contig ended after exon 6 and included a large insertion originating from downstream of the *para* gene.

A major motivation for pursuing PacBio sequencing following 10X genomics was that we did not identify a 10X contig that spanned the *para* gene. The 10X assembly did include *kdr* and a portion of the *para* gene in contig 253707 (6,808bp). However, contig 253707 ended at exon 6, which does not include sufficient flanking sequence to assist with *kdr* assay design. In contrast, the PacBio assembly generated a 31,996bp contig (tig00032677) that spanned the entire gene. Further analysis of the 10X contig versus the PacBio contig revealed a 671bp insertion in the 10X contig that maps in the reverse-complement direction downstream of the *para* gene (Figure 1).

The *para* gene has a total of eight exons and spans 11,104bp. The complete gene was not identified by the maker pipeline, but support for multiple exons from *Cx. quinquefasciatus* protein alignments were included in the output. To complete the annotations of the *para* gene coding regions in tig00032677, we used the *Cx. quinquefasciatus* supercont3.182 (contains *para* gene) and associated annotations as input for the Geneious annotation software. Using this approach, all eight exons were identified. Most of the exons had high percent similarity between *Cx. tarsalis* and *Cx. quinquefasciatus* (up to 99%), but exon 1 was more divergent with 62.4% similarity (Figure 1).

### Sex locus

In an attempt to annotate the sex locus in *Cx. tarsalis*, we utilized both the 10X genomics contigs and PacBio contigs to generate scaffold sequences near the sex locus on Chromosome 3. Molecular markers flanking the the sex locus have been previously described, including the microsatelite markers CUTB210 and CUTB218 that flank the sex locus by 1.5 cM and 1.4 cM, respectively (Venkatesan *et al*. 2009). CUTB210 (DQ682690) is a 553bp segment on the telemeric side of the sex locus and maps to a 17kb contig of CtarK1 at position tig00029712:10,209-10,765. CUTB210 has 9 additional “CT” tandem repeats at position 10,529 and 1 SNP at position 10,626 versus our CtarK1 reference. To scaffold contigs across the sex locus, we used a 215.5kb 10X contig (contig 430), which aligns with tig00029712 at position 430:123,826-135,535. Then, we identified the PacBio tig00005857 (113kb), which maps to the last 20kb of 10X contig 430. Contig 305071 (10X; 21,693bp) maps to approximately the last 16kb of tig00005857 (96,444-113,183), extending the scaffold approximately another 5kb. The final CUTB210-scaffold is 321,101bp.

The CUTB218 sequence is 481bp and resides on the centromeric side of the sex locus. This sequence failed to map to CtarK1. However, CUTB218 maps to the extreme end of the 10X contig-233502 (29kb) with approximately 20 more CT tandem repeats than the 10X reference sequence. Using 10X contig-233502 as a scaffold, we identified the PacBio tig00011961 (33kb), which maps to the first 16,575bp. Several large insertions were identified in 10x contig versus PacBio, including one 5kb stretch of N’s (supplemental figure 1). Another large PacBio contig (tig00000802; 164kb) mapped to the opposite end of contig-233502. Approximately 9.5kb overlapped at the end of this contig, however the last 2.5kb (which includes CUTB218) was either not in synteny or didn’t align at all (supplemental figure 2). Due to the poor alignment at the extreme end of contig-233502, including the CUTB218 sequence, we excluded tig00000802 from the CUTB218 scaffold. Thus, the final CUTB218 scaffold was 45,566bp. Attempts to scaffold using The *Cx. quinquefasciatus* genome did not improve these scaffolds. To facilitate future efforts to characterize the sex locus, we have included these sequences in Supplemental File 1.

Repetitive elements were annotated in parallel using RepeatMasker and a custom repeat library. In total, 60.8% of CtarK1 was annotated as a repeat feature. This is double the estimate from *Cx. quinquefasciatus* (Arensburger et al. 2010), indicating that the *Cx. tarsalis*-specific genome expansion (800Mb vs 579Mb) may be mostly composed of transposable elements.

### Phylogenetic analysis

Based on multiple-species alignments of complete mitochondrial genomes (∼15kb), we estimated the placement of *Cx. tarsalis* on a phylogenetic tree with other sequenced mosquito species and *Drosophila melanogaster* as an outgroup. The estimated divergence time between *Cx. tarsalis* and *Cx. quinquefasciatus* was 15.8-22.2 MYA (95% Credible Interval; Figure 3). The more recent split between *Culex tarsalis* and *Aedes mosquitoes* versus *Anopheles (Severson and Behura 2012)* and the divergence estimate between *An. gambiae* and *An. coluzzii* (95% Credible Interval = 0.07-3.45) was consistent with the previous estimate of 0.061 MYA (Thawornwattana *et al*. 2018).

**Figure 2.**
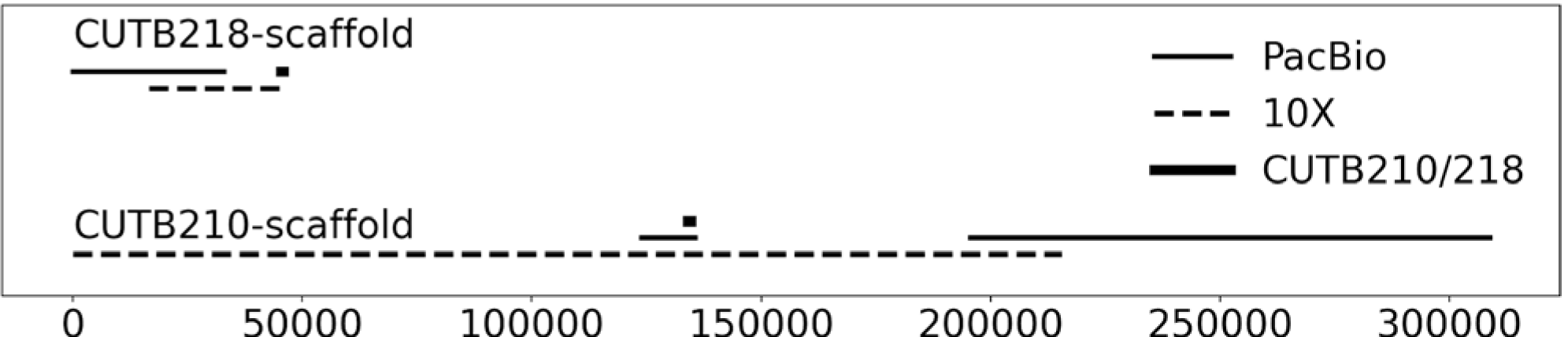
Scaffolding of 10X and PacBio contigs flanking the sex locus. In an attempt to sequence across the sex locus, 10X Genomics (dashed lines) and PacBio (solid lines) contigs were combined at the previously described markers (black squares) that flank the telomeric (CUTB210) and the centromeric side (CUTB218) of the sex locus (Venkatesan *et al*. 2009).

**Figure 3.**
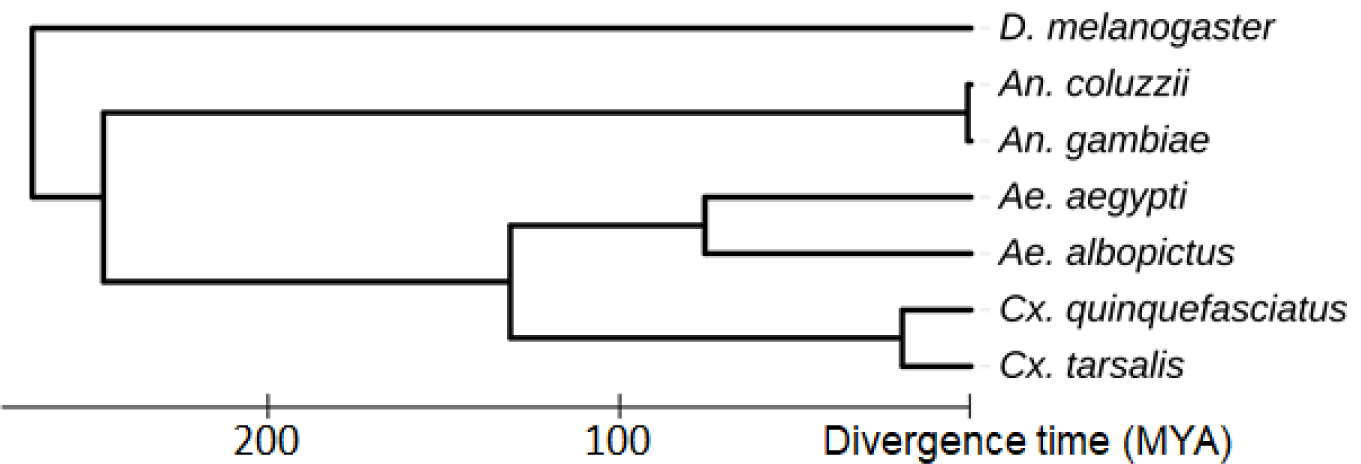
Mitochondrial phylogeny. This Bayesian phylogeny places *Cx. tarsalis* in the context of other sequenced mosquito species and *D. melanogaster* as an outgroup. This estimate was based on multiple mitochondrial sequence alignments (15,203bp) and calibration constraints for *Aedes* species, between *Culex* and *Aedes*, between *Culicinae* and *Anopheles*, and between *Drosophila* and *Culicidae* (Chen *et al*. 2015). The divergence time estimate between *Cx. tarsalis* and *Cx. quinquefasciatus* is 15.8-22.2 MYA.

### Conclusions

Here, we present the first genome assembly and genomic analysis for *Cx. tarsalis*. Based on full mitochondrial genome alignments, we estimate that *Cx. tarsalis* diverged from *Cx. quinquefasciatus* 15.8-22.2 MYA, well after the last common ancestor between *Aedes albopictus* and *Aedes aegypti*. This is an important first step toward understanding the genome evolution between these important vector species. This *Cx. tarsalis* genome assembly (790Mb) is 27% larger than the *Cx. quinquefasciatus* genome (578Mb), and this difference appears to be mostly composed of transposable elements. Future work will include comparative genomics within *Culex* versus *Anopheles* species (Neafsey *et al*. 2015), with a particular focus on genes involved in diapause. The sex locus has also not yet been characterized in any *Culex* species. Understanding this basic biology is important for understanding the evolution of sex determination in mosquitoes and may also lead to innovative new vector control strategies involving sex ratio distorters.

## Acknowledgements

We thank the Alameda County Mosquito Abatement District for sponsoring a portion of the genome assembly work, and especially Eric Haas-Stapleton and Ryan Clausnitzer for their enthusiastic support of our research on *Culex tarsalis* genomics. We acknowledge funding from an Innovative Development Award from the UC Davis Academic Senate and the Vector-Borne Disease Pilot Grant program of the UC Davis School of Veterinary Medicine. The sequencing was carried out by the DNA Technologies Core and Expression Analysis Core at the UC Davis Genome Center, supported by NIH Shared Instrumentation Grant 1S10OD010786-01. MM and CMB acknowledge funding support from the Pacific Southwest Center of Excellence in Vector-Borne Diseases funded by the U.S. Centers for Disease Control and Prevention (Cooperative Agreement 1U01CK000516). CTB was supported by the Gordon and Betty Moore Foundation under grant GBMF4551. JLR was supported by NIH grant 1R01AI150251.

## Data availability

The reference genome and annotation file were deposited at the open science framework DOI: 10.17605/OSF.IO/MDWQX

## References Cited

Arensburger, P., K. Megy, R. M. Waterhouse, J. Abrudan, P. Amedeo et al., 2010 Sequencing of Culex quinquefasciatus establishes a platform for mosquito comparative genomics. Science 330: 86–88.

Buth, J. L., R. A. Brust, and R. A. Ellis, 1990 Development time, oviposition activity and onset of diapause in Culex tarsalis, Culex restuans and Culiseta inornata in southern Manitoba. J. Am. Mosq. Control Assoc. 6: 55–63.

Chen, X.-G., X. Jiang, J. Gu, M. Xu, Y. Wu et al., 2015 Genome sequence of the Asian Tiger mosquito, Aedes albopictus, reveals insights into its biology, genetics, and evolution. Proc. Natl. Acad. Sci. U. S. A. 112: E5907–15.

Drummond, A. J., and A. Rambaut, 2007 BEAST: Bayesian evolutionary analysis by sampling trees. BMC Evol. Biol. 7: 214.

Edgar, R. C., 2004 MUSCLE: a multiple sequence alignment method with reduced time and space complexity. BMC Bioinformatics 5: 113.

Goddard, L. B., A. E. Roth, W. K. Reisen, and T. W. Scott, 2002 Vector competence of California mosquitoes for West Nile virus. Emerg. Infect. Dis. 8: 1385–1391.

Johnston, J. S., A. Bernardini, and C. E. Hjelmen, 2019 Genome Size Estimation and Quantitative Cytogenetics in Insects, pp. 15–26 in Insect Genomics: Methods and Protocols, edited by S. J. Brown and M. E. Pfrender. Springer New York, New York, NY.

Kramer, L. D., J. L. Hardy, S. B. Presser, and E. J. Houk, 1981 Dissemination barriers for western equine encephalomyelitis virus in Culex tarsalis infected after ingestion of low viral doses. Am. J. Trop. Med. Hyg. 30: 190–197.

Neafsey, D. E., R. M. Waterhouse, M. R. Abai, S. S. Aganezov, M. A. Alekseyev et al., 2015 Highly evolvable malaria vectors: The genomes of 16 Anopheles mosquitoes. Science 347: 1258522.

Reisen, W. K., 2013 Ecology of West Nile virus in North America. Viruses 5: 2079–2105.

Reisen, W. K., 1995 Effect of temperature on Culex tarsalis (Diptera: Culicidae) from the Coachella and San Joaquin Valleys of California. J. Med. Entomol. 32: 636–645.

Reisen, W. K., 1986 Overwintering Studies on Culex tarsalis (Diptera: Culicidae) in Kern County, California: Life Stages Sensitive to Diapause Induction Cues. Ann. Entomol. Soc. Am. 79: 674–676.

Reisen, W. K., 2012 The Contrasting Bionomics of Culex Mosquitoes in Western North America. J. Am. Mosq. Control Assoc. 28: 82–91.

Reisen, W. K., Y. Fang, and V. M. Martinez, 2014 Effects of temperature on the transmission of West Nile virus by Culex tarsalis (Diptera: Culicidae). J. Med. Entomol. 43: 309–317.

Reisen, W. K., H. D. Lothrop, and T. Thiemann, 2013 Host Selection Patterns of Culex tarsalis (Diptera: Culicidae) at Wetlands Near the Salton Sea, Coachella Valley, California, 1998–2002. J. Med. Entomol. 50: 1071–1076.

Reisen, W. K., P. T. Smith, and H. D. Lothrop, 1995 Short-term reproductive diapause by Culex tarsalis (Diptera: Culicidae) in the Coachella Valley of California. J. Med. Entomol. 32: 654–662.

Ribeiro, J. M. C., I. Martin-Martin, F. R. Moreira, K. A. Bernard, and E. Calvo, 2018 A deep insight into the male and female sialotranscriptome of adult Culex tarsalis mosquitoes. Insect Biochem. Mol. Biol. 95: 1–9.

Severson, D. W., and S. K. Behura, 2012 Mosquito genomics: progress and challenges. Annu. Rev. Entomol. 57: 143–166.

Spadoni, R. D., R. L. Nelson, and W. C. Reeves, 1974 Seasonal Occurrence, Egg Production, and Blood-feeding Activity of Autogenous Culex tarsalis,. Ann. Entomol. Soc. Am. 67: 895–902.

Thawornwattana, Y., D. Dalquen, and Z. Yang, 2018 Coalescent Analysis of Phylogenomic Data Confidently Resolves the Species Relationships in the Anopheles gambiae Species Complex. Mol. Biol. Evol. 35: 2512–2527.

Thiemann, T. C., D. A. Lemenager, S. Kluh, B. D. Carroll, H. D. Lothrop et al., 2012 Spatial variation in host feeding patterns of Culex tarsalis and the Culex pipiens complex (Diptera: Culicidae) in California. J. Med. Entomol. 49: 903–916.

Venkatesan, M., K. W. Broman, M. Sellers, and J. L. Rasgon, 2009 An initial linkage map of the West Nile Virus vector Culex tarsalis. Insect Mol. Biol. 18: 453–463.

Venkatesan, M., and J. L. Rasgon, 2010 Population genetic data suggest a role for mosquito-mediated dispersal of West Nile virus across the western United States. Mol. Ecol. 19: 1573–1584.

Waterhouse, R. M., M. Seppey, F. A. Simão, M. Manni, P. Ioannidis et al., 2018 BUSCO Applications from Quality Assessments to Gene Prediction and Phylogenomics. Mol. Biol. Evol. 35: 543–548.

Zdobnov, E. M., F. Tegenfeldt, D. Kuznetsov, R. M. Waterhouse, F. A. Simão et al., 2017 OrthoDB v9.1: cataloging evolutionary and functional annotations for animal, fungal, plant, archaeal, bacterial and viral orthologs. Nucleic Acids Res. 45: D744–D749.

Ziegler, R., S. Whyard, A. E. R. Downe, G. R. Wyatt, and V. K. Walker, 1987 General esterase, malathion carboxylesterase, and malathion resistance in Culex tarsalis. Pestic. Biochem. Physiol. 28: 279–285.

